# Activation of specific mushroom body output neurons inhibits proboscis extension and feeding behavior

**DOI:** 10.1101/768960

**Authors:** Justine Chia, Kristin Scott

## Abstract

The ability to modify behavior based on prior experience is essential to an animal’s survival. For example, animals may become attracted to a previously neutral odor or reject a previously appetitive food source upon learning. In *Drosophila*, the mushroom bodies (MBs) are critical for olfactory associative learning and conditioned taste aversion, but how the output of the MBs affects specific behavioral responses is unresolved. In conditioned taste aversion, *Drosophila* shows a specific behavioral change upon learning: proboscis extension to sugar is reduced after a sugar stimulus is paired with an aversive stimulus. While studies have identified MB output neurons (MBONs) that drive approach or avoidance behavior, whether the same MBONs impact proboscis extension behavior is unknown. Here, we tested the role of MB pathways in modulating proboscis extension and identified 10 *MBON split-GAL4* lines that upon activation significantly decreased proboscis extension to sugar. Activating several of these lines also decreased sugar consumption, revealing that these MBONs have a general role in modifying feeding behavior beyond proboscis extension. Although the MBONs that decreased proboscis extension and ingestion are different from those that drive avoidance behavior in another context, the diversity of their arborizations demonstrates that a distributed network influences proboscis extension behavior. These studies provide insight into how the MB flexibly alters the response to taste compounds and modifies feeding decisions.

## Introduction

A key role of the brain is to prioritize relevant sensory information to guide behavior. Animals exhibit innate behaviors to a variety of sensory stimuli including tastes and odors, and the ability to modify those behaviors based on contextual cues and prior experience is essential to an animal’s survival.

In *Drosophila*, the mushroom body (MB) has long been implicated as a center for learning and memory, and has been studied most extensively in the context of olfactory associative learning (1–4). The dendrites of the principal cells of the MB, Kenyon cells (KCs), receive sparse, random synaptic inputs from olfactory projection neurons (5). The parallel axonal fibers of the KCs form the MB lobes, the output region of the MB. The axonal lobes comprised of ∼2000 KC axons are beautifully organized into 15 compartments, defined anatomically by the dendrites of 21 types of MB output neurons (MBONs) (6). These compartments also contain the axon terminals of 20 types of dopaminergic neurons (DANs), which similarly tile the MB lobes. The DANs convey signals of reward or punishment for sensory associations (7-14). In Pavlovian terminology for a classical conditioning paradigm, the odor is designated the conditioned stimulus (CS), and the DANs carry the unconditioned stimulus (US).

In the current model of olfactory associative learning, behavior is determined through the summation of activity in different MB compartments. Some compartments encode approach behavior, and others encode avoidance (15). In a naïve animal, the total valence carried by the positive and negative compartments is in balance, and thus there is no observed approach or avoidance behavior. During olfactory aversive learning, the balance tips towards avoidance behavior: long-term depression is thought to decrease the output in a compartment that signals positive valence (16). Using similar logic, after appetitive olfactory learning, DANs carrying a rewarding signal decrease the activity in aversive MBONs, leading to increased acceptance. The sum of synaptic changes drives the overall behavior toward avoidance or acceptance, thereby modifying the innate behavior.

In support of the model that individual MBONs encode a positive or negative valence, the behavioral roles of MBONs have been investigated through direct activation (15, 17). Optogenetic activation of some MBONs causes approach behavior, while activation of other MBONs causes avoidance behavior (15). Activation of other MBON subsets causes neither approach nor avoidance. A different subset of MBONs has been found to play a critical role in odor-seeking behavior (18). Although these studies argue that diverse MBONs can influence behavior, it is unclear whether MBONs signal a positive or negative valence in the context of multiple behaviors or how MBONs impact feeding behaviors like proboscis extension.

How do the outputs of the MBs influence specific behaviors? One possibility is that MB outputs alter the probability of response to a given sensory stimulus, promoting or inhibiting the response, independent of the nature of the response. In this model, MB outputs would serve to gate the probability of actions but not participate in action selection. One prediction of this model would be that the MBONs that signal a positive valence and promote approach might promote other behavioral responses as well. An alternative model is that MBON activity has behavioral specificity. For example, the MBONs that promote approach might be different from the MBONs that promote ingestion. To begin to examine this question, we tested the role of MB pathways in modulating an innate behavior, proboscis extension to sucrose.

The fly gustatory system is an excellent model to study how MB pathways modulate innate behaviors, as feeding decisions may be altered by learned associations, and importantly, there is a clear behavioral readout: the proboscis extension response (PER). For example, during conditioned taste aversion, a paired application of sugar to the tarsi and bitter to the proboscis results in a reduction of PER to sugar alone (19–22). During conditioned aversion, taste information is transmitted from the subesophageal zone (SEZ) to the MBs for learned associations (22–24). While studies have found that some components of the MBs are required for taste memory formation (22, 24, 25), how MBONs impact innate proboscis extension behavior has not been resolved.

In this study, we test the role of MB pathways in modulating PER. We find that a subset of MBONs drives inhibition of proboscis extension. Specifically, we identified 10 *MBON split-Gal4* lines that upon activation significantly decreased proboscis extension to sugar. Inhibiting neural activity in these *MBON split-Gal4* lines did not reciprocally regulate proboscis extension. Activating several of the identified *MBON split-Gal4* lines also decreased sugar consumption, revealing that these MBONs have a more general role in the feeding circuit beyond the proboscis extension motor program. In addition, activating dopaminergic inputs in 3 MB compartments also suppressed proboscis extension to sugar. The MBONs that decrease proboscis extension and ingestion are different from those mediating avoidance in another context. The diversity of MBON compartments that influences PER demonstrates that a distributed network influences behavior and is consistent with the role of MBs in generating context-dependent behavioral biases.

## Results

### Activation of MBONs suppresses proboscis extension

To identify the MB outputs that modulate proboscis extension, we screened 35 previously described *split-Gal4* lines that cover the 21 MBON types (15). We used the red-light-activated ion channel CsChrimson to optogenetically activate *split-Gal4* lines. Flies were raised on standard cornmeal food (controls) or all-trans-retinal-supplemented food (experimental animals) were starved for 1 day before being tested for proboscis extension. Proboscis extension was tested in response to simultaneous 635 nm light illumination to activate MBONs and 100 mM sucrose presentation to the tarsi. For 12 lines, we found that activation with CsChrimson caused a significant decrease in proboscis extension to sucrose (Fig 1A). One line (MB323B) had motor defects and leg folding upon activation and was excluded from further study.

**Fig 1.**
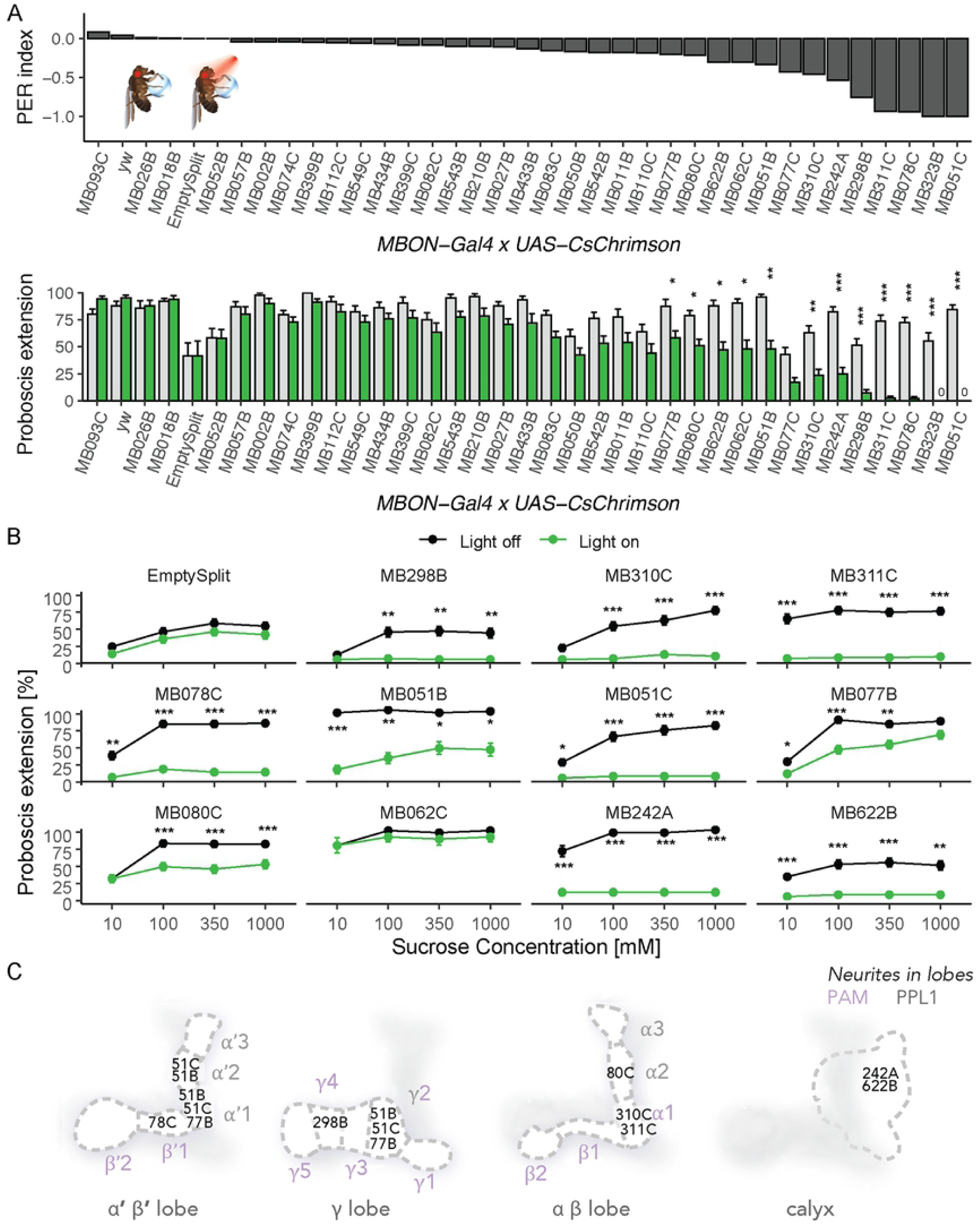
Identification of MBONs that suppress proboscis extension to sucrose. A) Behavioral screen for flies that change their rate of proboscis extension when MBONs are activated. *MBON split-Gal4* lines were crossed to *UAS-CsChrimson* for light induced activation and tested for proboscis extension to simultaneous red light and sucrose presentation to the tarsi. Extension rates were compared between flies fed retinal and those not. (n = 20-48 flies per line). Inset: Illustration of the PER assay. Top: results of the screen, ordered by PER index to reveal Gal4 lines with the greatest change in PER upon MBON activation. PER index = (Retinal – no retinal) / (Retinal + no retinal). Bottom: Same data ordered as in top, shown as mean ± SEM. Statistical significance was calculated using Wilcoxon Rank Sum tests (retinal versus no retinal) with Bonferroni correction for multiple comparisons, *p<0.05, **p < 0.01, ***p < 0.001. Green bars represent flies fed retinal. Grey bars represent flies not fed retinal. B) Retest of candidates causing the greatest PER suppression upon MBON activation with CsChrimson. Values represent mean (± SEM) fraction of flies presented with sucrose (black lines) and flies presented with sucrose and red light (green lines) exhibiting PER to the indicated concentrations of sucrose (n = 26-57). Asterisks denote statistically significant differences between flies in light and dark conditions. Statistical significance was calculated using a two-way ANOVA with Bonferroni correction, where *p<0.05, **p < 0.01, ***p < 0.001. C) Schematic of dendritic arborizations of the *MBON split-Gal4* lines that caused the greatest PER suppression phenotypes in the activation screen. Each lobe of the MB, as well as the calyx, is drawn separately for visual clarity. The name of each *MBON split-Gal4* is spatially localized to the compartments where it has dendritic arborizations. Colors indicate cluster of origin for DANs.

The 11 lines showing reduced proboscis extension upon MBON activation were re-tested for proboscis extension to a range of sucrose concentrations (10, 100, 350, 1000 mM) with and without CsChrimson activation of MBONs. Ten of the 11 lines showed significantly reduced proboscis extension to several sucrose concentrations upon CsChrimson activation (Fig 1B). One line (MB062C) did not show a phenotype upon retesting and was excluded from further study (Fig 1B). Importantly, the decrease in proboscis extension was not due to fly paralysis, as determined by measuring walking speed in a locomotor assay (Fig S1). The 10 *MBON split-Gal4* lines that reduced proboscis extension are not localized to any single compartment or lobe. Instead, their neurites arborize in 7 of the 15 mushroom body compartments and the calyx and belong to 7 different MBON cell-types: γ4>γ1γ2, α1, β’1, γ2α’1, α’2, α2sc, and the calyx (Fig 1C).

### Inhibition of MBON candidates does not influence proboscis extension

As activation of several *MBON split-Gal4* lines decreased proboscis extension to sucrose, we hypothesized that reducing neural activity in those MBONs would cause a reciprocal increase in PER. To test this, we expressed a temperature sensitive, dominant negative dynamin, Shibire^*ts*,^ using a transgene that drives strong expression (*20xUAS-Shi*^*ts1*^) in the 10 *MBON split-Gal4* lines that influenced proboscis extension. Flies were stimulated with 100 mM sucrose applied to the proboscis at 30–32°C to inhibit vesicle reuptake or at ∼22°C as same genotype controls. Flies tested at 30–32°C were pre-incubated for 15 minutes prior to the start of the experiment on a 30– 32°C heating block. Only 1 line, MB078C, showed increased PER at the restrictive temperature (Fig 2A).

**Fig 2.**
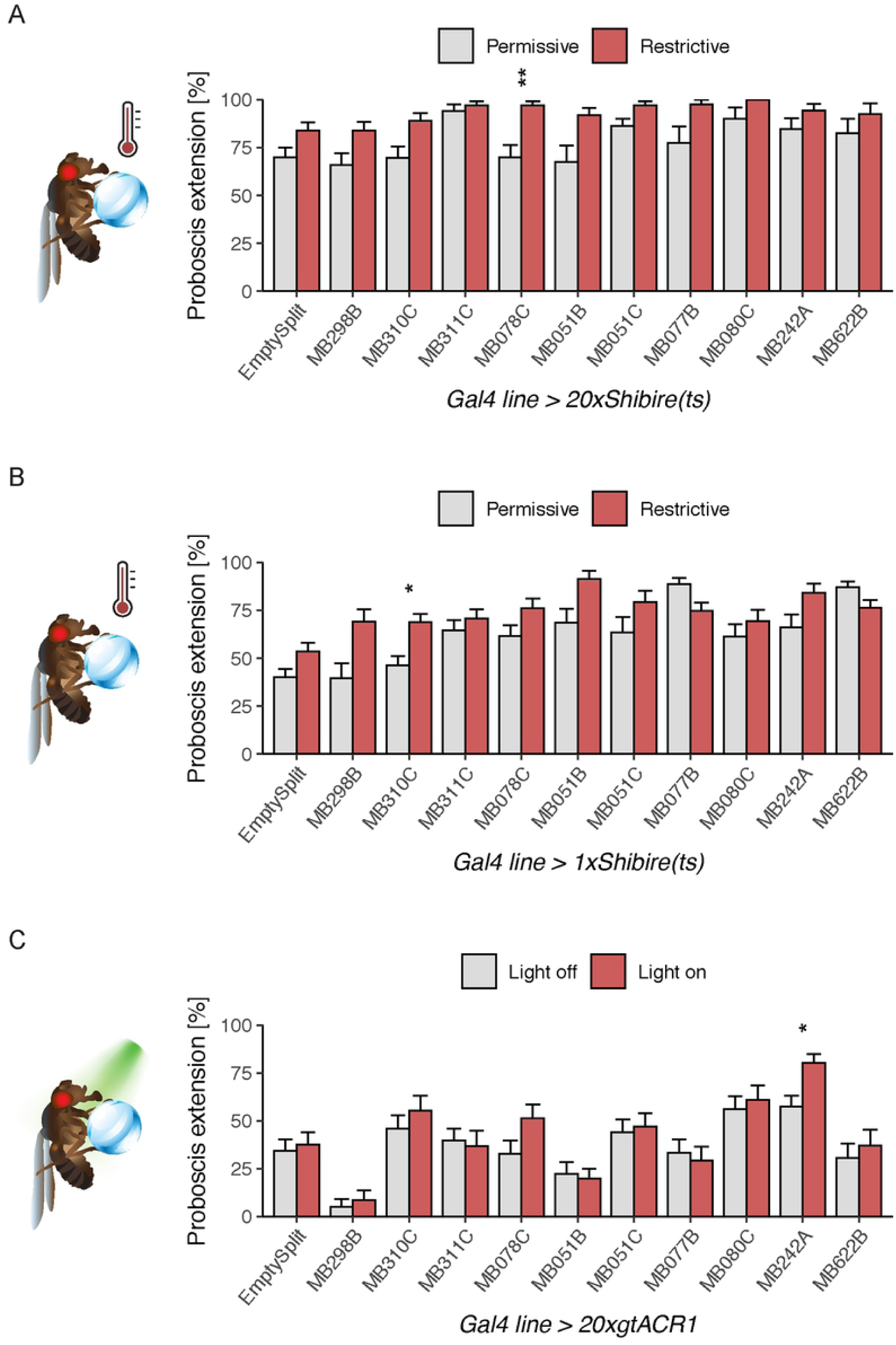
Decreasing neural activity in MBONs that suppress PER when activated has modest effects. A) MBON lines identified in our activation screen were conditionally silenced with 20xShibire^ts^ and PER to tarsal sugar presentation (100 mM sucrose) was recorded. Silencing with this method caused an increase in one line (MB078C). n = 18-59. Permissive temperature= 20-22°C, restrictive = 30–32°C. B) MBON candidates were conditionally silenced with 1xShibire^ts^ and PER to 50 mM sucrose on the legs was recorded. Silencing MBONs with this method caused an increase in PER rate in one line (MB310C). n = 20-74. C) Candidates were silenced acutely with the light-gated anion channelrhodopsin 20xgtACR1 and PER to 10 mM sucrose on the legs was recorded. Silencing MBONs with this method increased PER to sucrose in one line (MB242A). n = 16-36. For all graphs, error bars indicate mean ± SEM. Statistical significance was determined by Wilcoxon Rank Sum tests with Bonferroni correction for multiple comparisons, *p<0.05, **p<0.01.

Because the strong Shibire effector has been reported to produce phenotypes at the permissive temperature (15), we repeated these experiments using the weaker *1xUAS-Shi*^*ts1*^. We also altered the behavioral paradigm to stimulate with 50 mM sucrose instead of 100 mM sucrose, as PER to 100 mM sucrose under control conditions was high, creating the possibility of ceiling effects. Under these conditions, 1 of the 10 *MBON split-Gal4* lines (MB310C) showed increased proboscis extension upon neural silencing (Fig 2B).

Finally, we tested an additional acute silencing strategy that provides rapid light-triggered hyperpolarization. The light-gated anion channelrhodopsin, gtACR1, was expressed in candidate MBON lines. Flies were stimulated with 10 mM sucrose, as this concentration produced ∼50% PER in genotype controls. For each MBON line, the same genotype was examined in the presence of 635 nm light for neural silencing or under control conditions. We found 1 line (MB242A) where acute silencing with gtACR1 increased proboscis extension (Fig 2C).

Taken together, the neural silencing experiments argue that the *MBON split-Gal4* lines that inhibited proboscis extension when activated do not consistently alter proboscis extension when inhibited. One explanation may be that proboscis extension to sugar is modulated by MBONs although they are not a required component of the sensorimotor circuit. Instead, the proboscis extension motor program may be controlled by local SEZ circuits and an alternative pathway may relay taste information to the higher brain for learned associations. Alternatively, MBONs may not be intrinsically active and blocking activity in neurons that are already silent may not produce a phenotype. Another possibility is that multiple MBONs may need to be silenced in order to produce a phenotype (15). Regardless, these studies demonstrate that inhibiting single classes of MBONs does not strongly influence proboscis extension to sucrose.

### MBON activation also affects sugar consumption

We next asked whether MBONs have a more general role in influencing feeding behavior beyond the simple proboscis extension motor program. To address this, we investigated the effect of *MBON split-Gal4* line activation on sucrose consumption. We hypothesized that the set of MBONs whose activation suppresses proboscis extension would also decrease sucrose consumption. In flies starved 24 hours, we measured consumption of 100mM sucrose while activating the 10 *MBON split-Gal4* lines with a PER suppression phenotype using the red-light gated channel 10xChrimson88. Comparing consumption in the presence of red light (for activation) to consumption without red light (controls), we found that 3 lines (MB078C, MB311C, and MB242A) consumed less sugar upon activation (Fig 3). These MBONs provide inputs to the β’1 compartment, the α1 compartment and the MB calyx, respectively.

**Fig 3.**
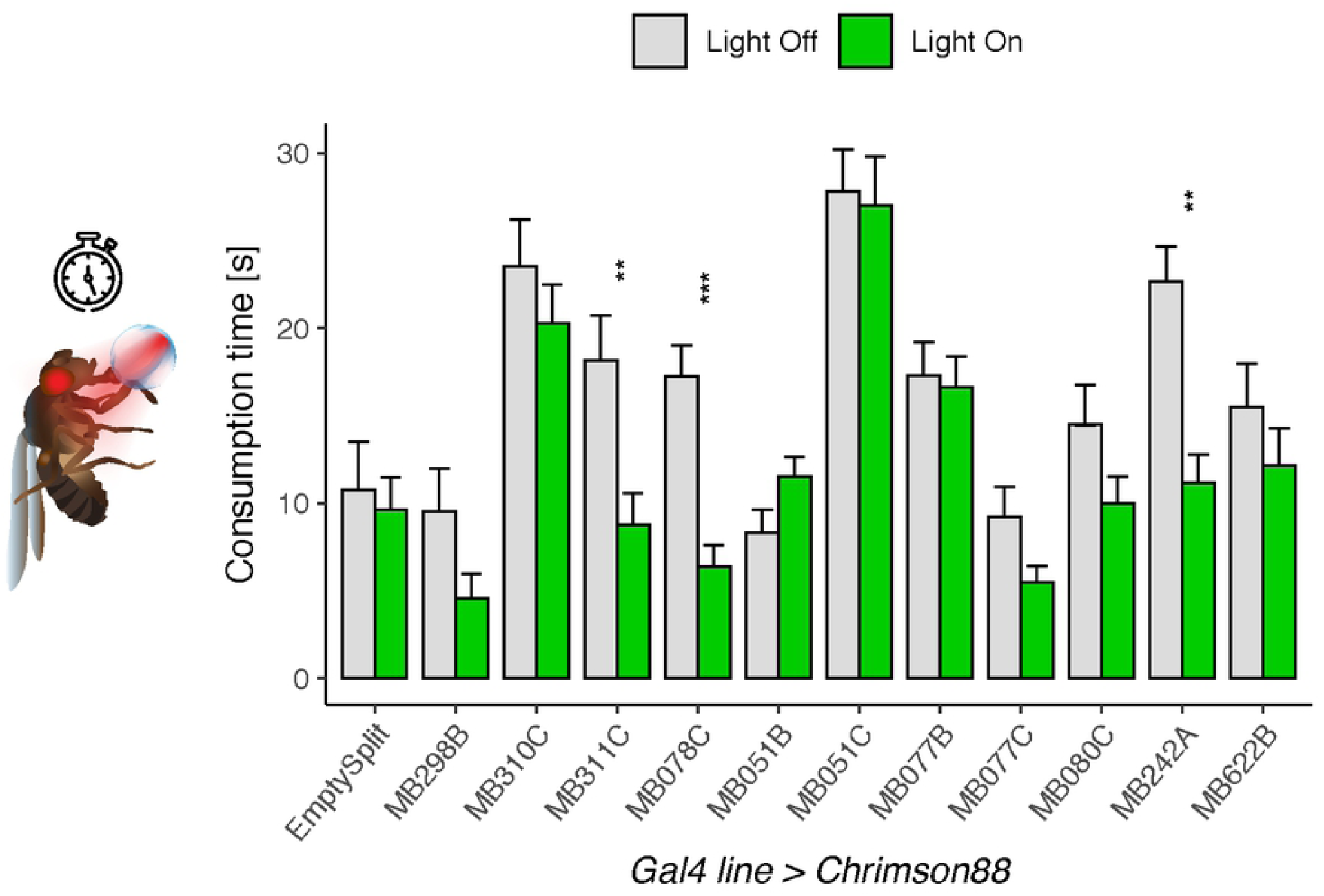
Activation of MBONs causes a reduction in sucrose consumption. Activation of 3 *MBON split-Gal4* lines (MB311C, MB078C, MB242A) using Chrimson88 caused decreased consumption of 100 mM sucrose. Values represent mean ± SEM. Statistical significance was calculated using unpaired Wilcoxon Rank Sum tests (light versus no light) with Bonferroni correction for multiple comparisons; **p<0.01, ***p<0.001; n = 15-42 flies per line.

### Screen for MB DANs regulating PER

In the current model of the MB, activity is balanced between different compartments to drive overall behavior. The dopaminergic inputs to MB compartments are thought to change the strength of synaptic connections between MB neurons and MBONs, mainly through their inputs onto the KCs conveying sensory information (15, 26, 27). Whether those synaptic connections are weakened or strengthened appears to be context-dependent and compartment-specific: in some studies DANs inhibit MBON outputs through long term depression (LTD) (16, 17, 28–31), while in other studies DANs have been reported to potentiate KC to MBON connections (17, 32– 34). The timing of the DAN and sensory inputs is critically important for determining synaptic potentiation or depression (16, 35).

To investigate whether specific DANs modulate PER and whether they are associated with the same compartments innervated by MBONs that influence PER, we conducted an unbiased screen of DANs. Since activating MBONs innervating γ4>γ1γ2, α1, β’1, γ2α’1, α’2, α2sc, and the calyx suppressed PER, we hypothesized that the DANs innervating those same compartments would cause an increase or decrease in PER.

To conduct the screen, we crossed 33 *DAN split-Gal4* lines to *10x UAS-Chrimson88,* and recorded proboscis extension to sucrose. Flies starved 24 hours were stimulated with 10 mM sucrose presented to the tarsi in the absence (controls) or presence of red light to activate DANs and tested for proboscis extension. This sucrose concentration was chosen to provide a dynamic range capable of detecting increases or decreases in PER rates. Upon activation with red light, 4 lines showed decreased PER to sucrose (Fig 4). Activation of one line (MB438B) caused legs to fold in the red light condition and was thus excluded from analysis. The remaining DAN lines causing decreased PER mainly innervate the PAM-β’2(amp) and PAM-α1 compartments (as well as weakly PAM-γ5 and PAM-β1).

**Fig 4.**
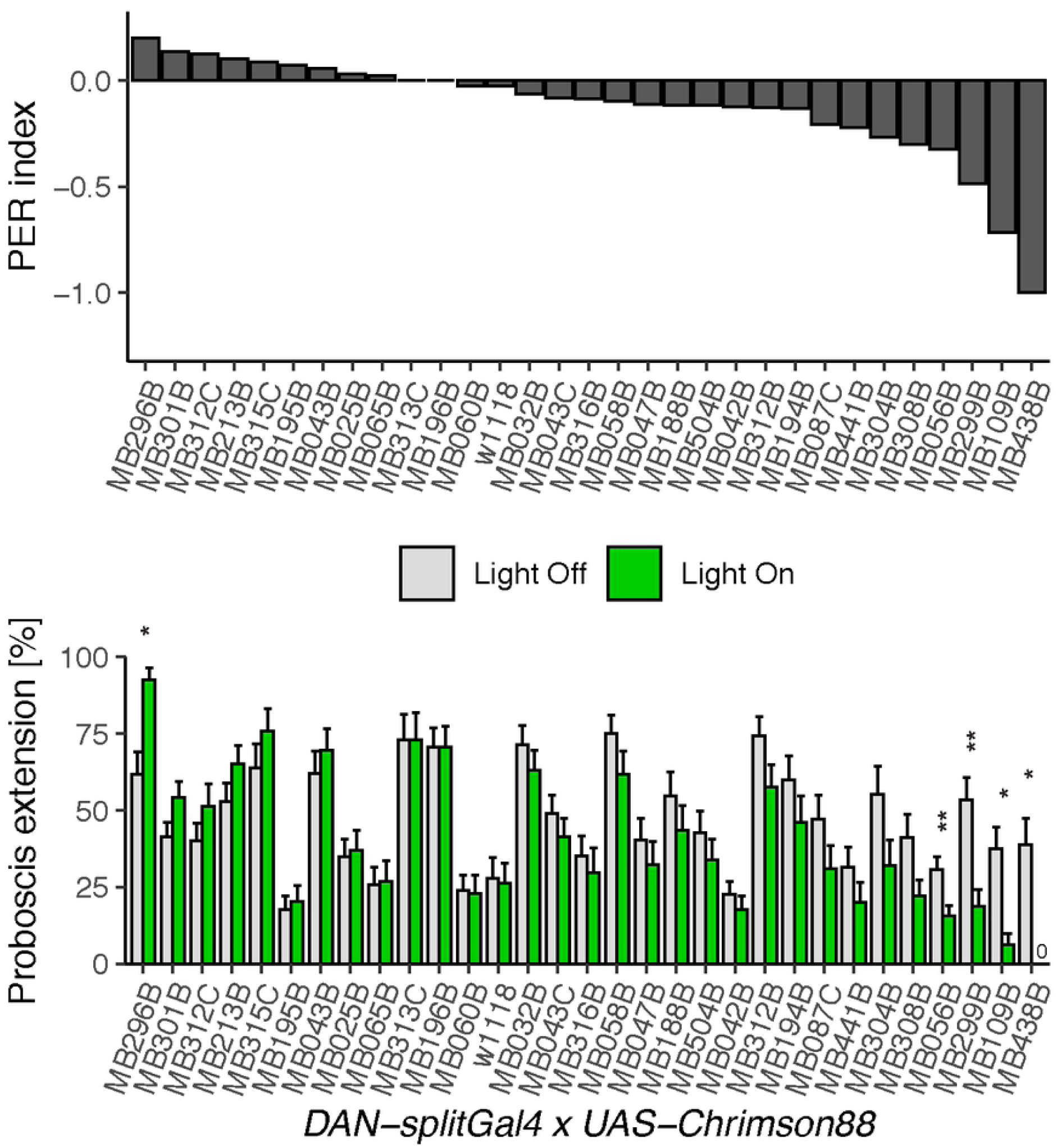
Identification of DANs that modulate PER to sucrose. Behavioral screen for flies that change proboscis extension rate when DANs are activated. *DAN split-Gal4* lines were crossed to *UAS-Chrimson88* for light induced activation and tested for proboscis extension to 10 mM sucrose presentation to the tarsi, and then simultaneous sucrose presentation to the tarsi and red laser light. Extension rates were compared between light and dark conditions in the same fly (n = 24-81 flies per line). Values represent mean ± SEM. Statistical significance was calculated using paired Wilcoxon Rank Sum tests (light versus no light) with Bonferroni correction for multiple comparisons, *p < 0.05, **p < 0.01.

In addition to these lines whose activation decreased PER, we also found one *DAN split-Gal4* line whose activation caused spontaneous PER: MB296B, a *split-Gal4* for PPL-γ2α’1 (Fig. 4). We re-tested this line with the effector CsChrimson and found robust PER to red light (Fig. S2A). Chronic silencing of these neurons with the inward rectifying potassium channel Kir2.1 resulted in increased PER compared to genetic controls (Fig S2B). Acute silencing with gtACR1 did not have a significant effect (Fig S2C). MB296B labels some neurons outside PPL-γ2α’1 in the SEZ where gustatory sensory axons terminate and proboscis extension motor neurons are located (Fig. S2D). To address the contribution of non-PPL-γ2α’1 in MB296B, we used an intersectional strategy to restrict CsChrimson expression to the SEZ using a Hox gene promoter that overlaps with the expression of MB296B (Fig. S2D). In flies that express the red-light activated channel in the SEZ neurons of MB296B, there was PER to red light in some flies (Fig. S2E), suggesting that the SEZ neurons are responsible for some of the PER phenotype. Since red light induced PER in only a small fraction of flies expressing CsChrimson in the SEZ neurons, it is possible that a combination of the SEZ neurons and the PPL neurons contributes to proboscis extension.

## Discussion

In this study, we investigated the MBONs that influence proboscis extension and consumption behaviors. Pairing a sucrose stimulus with optogenetic activation of MBONs revealed that 10 *MBON split-Gal4* lines reliably decreased PER to sucrose. In addition, 3 lines linked to decreased PER also decreased sucrose consumption upon activation. None of the MBONs with activation phenotypes showed consistent reciprocal phenotypes upon decreasing neural activity. We also examined whether activation of DAN inputs into the MBs influenced PER to sucrose and identified 3 lines that decreased PER (Fig 5). These studies provide insight into how MBONs influence taste responses and encode behavior.

**Fig 5.**
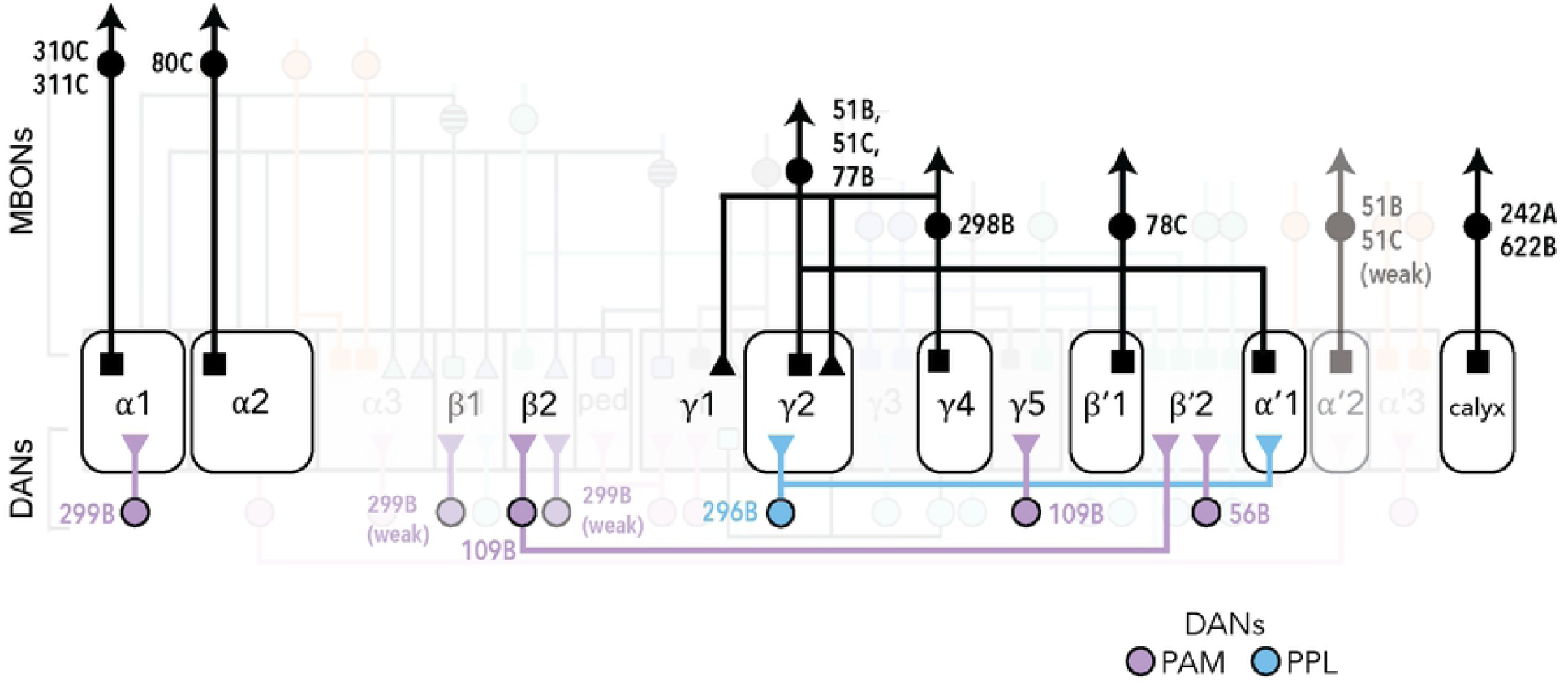
Schematic summary of MBONs and DANs whose activity influence PER. Schematic of the MB circuits of DAN inputs (bottom) and MBON outputs (top) whose activity influences PER. The 7 MB lobe compartments where MBON activation suppresses PER are shown in the grey rectangles (middle). Dendritic arborizations are represented by squares; triangles represent axonal arborizations. The cluster of origin of the DANs is indicated by the color of the label (purple for PAM and blue for PPL). MB051B, MB051C, and MB077B contain dendrites in 2 compartments. The α1 compartment is the one MB compartment whose activity consistently influenced feeding behavior (both PER and consumption). This diagram was modified from Figure 17 in (28).

The current model of MBON function proposes that each MBON has a positive or negative valence and the combined activity of MBONs determines whether the stimulus is attractive or aversive. Consistent with this model, previous studies have identified MBONs that drive approach and others that drive avoidance. Here, we tested MBONs to see whether they affect PER to sucrose and found that several *MBON split-Gal4* lines (MB298B, MB310C, MB311C, MB078C, MB051B, MB051C, MB077B, MB080C, MB242A, and MB622B) decreased PER. Of these, optogenetic activation studies have implicated MB298B in innate avoidance and MB077B in innate approach (28). The other MBONs that decreased PER did not show approach or avoidance behavior upon optogenetic activation. MB077B has also been shown to be required for food and odor seeking behavior in hungry flies (18), and silencing MB310C increases food seeking behavior (18). Additionally, MB080C has been previously described as having a PER suppression phenotype upon activation (22); the other lines identified here were not tested.

The observation that the majority of MBONs that suppress PER are different from those that cause innate avoidance or decrease yeast seeking argues against a simple model where MBONs have a fixed valence and activity is additive. Instead, the finding that the MBONs for PER inhibition are different from those that drive avoidance or decreased seeking suggests two different models for MBON function. One model is that MBONs have behavioral specificity, such that there are different MBONS for PER versus avoidance. An alternative model is that the context in which MBONs are activated determines their contribution to behavior. In our studies, flies were starved for 24 hours and were presented with sucrose. Both starvation and sucrose detection influence activity in MBs and may alter net MBON activity and the contributions of specific MBONs to behavior (9). Thus, our results do not distinguish whether different MBONs contribute to different behaviors or whether different MBONs have different weights in different contexts. Future studies are warranted to distinguish these models. However, given the number of MBONs that suppress PER and the number of compartments they innervate, the behavioral specificity model seems unlikely. Instead, these results are consistent with a more nuanced picture in which the role of MBONs in influencing behavior is context-dependent. Importantly, our studies argue against the notion that MBONs that drive aversion in one context are universally aversive in all contexts.

The finding that many MBONs reduce the probability of proboscis extension is consistent with the view that the MB is a complex, distributed circuit. The lines with the greatest PER suppression phenotype cover 7 different cell types: γ4>γ1γ2, α1, β’1, γ2α’1, α’2, α2sc, and the calyx, demonstrating that multiple compartments can influence this behavior (Fig. 5). In addition, 3 dopaminergic inputs into the MBs were also sufficient to regulate PER. These innervate the PAM-β’2(amp) and PAM-α1 compartments as well as weakly label PAM-γ5 and PAM-β1. There is no clear correlation between the DAN and MBON compartments that inhibit PER, as only the α1 compartment contains DANs and MBONs decreasing PER. Although the DAN activation experiments demonstrate that MB inputs are also sufficient to influence PER, how DAN activation propagates to MBONs to regulate behavior in these experiments is unresolved. It is possible that simultaneous activation of multiple DANs would elicit stronger effects on behavior.

One caveat with these studies is that artificial activation is not physiological and tests the case of strong activation. Under physiological conditions, there are likely more nuanced dynamics that drive behavior. Still, we find strong evidence that multiple MB compartments are able to modulate proboscis extension and feeding behavior.

Unlike activation, conditional silencing of MBONs had modest effects on PER in our study, suggesting many parallel pathways or compensatory mechanisms. One possibility is that proboscis extension is mediated by local SEZ circuits and modulated by MBONs. In this scenario, PER circuits may be influenced by MBONs but MBONs do not normally contribute to the behavior. Alternatively, multiple MBONs may contribute to the behaviors, such that silencing a large number of MBONs may be necessary to see a strong phenotype.

Overall, this work investigated how the activity of MB inputs and outputs impinges on feeding behavior by characterizing mushroom body neurons that impact proboscis extension to sucrose. We demonstrate that activation of several MBONs and DANs influences proboscis extension to sucrose, consistent with the view that a distributed network of MB compartments modulates behavior.

## Materials and Methods

### Fly stocks

The following fly lines were used: *MBON split-Gal4* lines (28);n*20xUAS-CsChrimson-mVenus* (attP18) (36); *10xUAS-Chrimson88-tdTomato3.1* (attp18) (David Anderson lab); *UAS-Shibire*^*ts*^ (37); *20xUAS-Shibire*^*ts*^ (38); *20xUAS-gtACR1* (39); *UAS-Kir2.*1 (40); *DAN split-Gal4* lines (28); *w*^*1118*^; pBPp65ADZpUw (attP40); pBPZpGAL4DBDUw (attP2) (Empty Split) (41); *20xUAS-dsFRT-CsChrimson-mVenus (42)*; *LexAop-FLP (43)*; *scr-LexA (44).*

*Drosophila* stocks were maintained at 25°C except those containing temperature-sensitive transgenes (*Shibire*^*ts*^) which were raised at 19°C. Flies were reared on standard fly food, except in experiments involving CsChrimson, Chrimson88, and gtACR1, which were transferred to food containing 0.4mM retinal prior to experiments.

### Behavior

Mated female flies, 4-9 days post-eclosion, were used for behavioral studies. Animals were starved for 20-26 hours in a vial with a kimwipe wet with 3 mL double distilled water.

For optogenetic experiments, flies with *UAS-Chrimson88, UAS-CsChrimson*, or *UAS-gtACR1* transgenes were raised in the dark at 25°C. Two days prior to testing, flies were transferred to fresh food with 0.4 mM all-trans retinal (Sigma). Flies were then starved on a wet kimwipe with the same retinal concentration. For activation experiments (CsChrimson, Chrimson88), flies were assayed one at a time with 635 nm light (LaserGlow). Silencing with gtACR1 was done using a custom-built LED panel (530nm) or 530-535nm laser (LaserGlow). For 1xShibire^ts^ and 20xShibire^ts^ experiments, flies were reared at either 19°C or 23°C. During the experiment, mounted flies were incubated at 30–32°C on a heating block for 15 minutes prior to the start of and throughout the experiment. For Kir2.1 experiments, flies were reared at 25°C.

#### Proboscis Extension Response Assay

PER was performed as described (45), except that each animal was treated as an independent data point. Briefly, flies were mounted on glass microscope slides with nail polish or UV glue (12 flies /slide for optogenetic experiments, and 36 flies/ slide for Kir2.1 and Shibire experiments), and allowed to recover for 15 minutes before being placed in a dark, humidified chamber for 2-5 hours. Prior to testing and between trials, flies were allowed to drink water ad libitum.

In the CsChrimson screen, we simultaneously activated MBONs while presenting 100 mM sucrose to the tarsi. This concentration was chosen because it is a moderately appetitive stimulus that results in proboscis extension ∼50% of the time in control flies. Flies were water-satiated before the experiment and between trials, and presented with the tastant and red light until proboscis extension was observed, for up to 5 seconds. Flies were given a score of 0 (for no extension) or 1 (for full extension), and the average was taken across two trials.

For thermogenetic silencing experiments (20xShibire^ts^ and 1xShibire^ts^), flies were assayed with 100 mM sucrose and 50 mM sucrose, respectively. For Kir2.1 experiments, flies were presented 30 mM sucrose.

For subsequent optogenetic (Chrimson88, gtACR1) PER assays with both MBONs and DANs, flies were assayed using 10 mM sucrose because 50 mM sucrose elicited close to 100% PER in the dark condition for some lines.

#### Temporal Consumption Assay

Temporal consumption assays were performed as described (23). Flies were glued onto glass slides using nail polish or UV glue, then allowed to recover in a humidified chamber for 2-4 hours. Each fly was water-satiated, then presented with 100 mM sucrose on the proboscis and forelegs. Cumulative drinking time over 10 consecutive presentations was recorded.

#### Locomotor Assay

Flies were gently aspirated into a circular bowl chamber made of 1.5% agarose, 44 mm in diameter [Bidaye et al., unpublished]. The stimulation protocol was 60s off, 30s pulsing 633nm light at 50 Hz. Light was delivered using a custom LED panel. Freely moving flies were videotaped under IR illumination using a Blackfly camera. The movie was subsequently analyzed using the Ctrax software suite version 0.3.9 (46). The total distance walked was computed and used to generate a mean distance traveled for each genotype assayed.

### Immunohistochemistry

Dissection and immunohistochemistry of 9- to 14- day old female fly brains were performed using the Janelia *split-Gal4* screen protocol (https://www.janelia.org/project-team/flylight/protocols) with small modifications: the incubation time for both primary and secondary antibodies was 3-7 days. Primary antibodies were chicken anti-GFP (Life Technologies, 1:500) and mouse anti-nc82 (Developmental Studies Hybridoma Bank, Iowa City, IA 1:500). Secondary antibodies were (both Invitrogen at 1:100): 488 anti-chicken, 647 antimouse. Images were acquired on a Zeiss confocal microscope. Brightness and contrast were adjusted using FIJI.

### Statistical Analyses

Statistical analyses were done using R (ggPubR). For PER in response to red light and sucrose in the CsChrimson screen, Wilcoxon Rank Sum tests with Bonferroni correction for multiple comparisons were used to compare the PER rate in retinal-exposed vs. no-retinal animals. For all other optogenetic PER assays, paired Wilcoxon tests with Bonferroni correction were used, since individual animals were tested both in the dark and light conditions. For PER experiments with Shi^ts1^ unpaired Wilcoxon tests were used since different populations of genetically identical animals were tested at the permissive temperature (22°) and the restrictive temperature (32°). For TCA experiments, Wilcoxon Rank Sum tests were used with Bonferroni correction for multiple comparisons.

## Acknowledgments

We thank members of the Scott lab for discussions and advice. This research was supported by a grant from the NIDCD 1R01DC013280 (KS) and an NSF predoctoral fellowship (JC).

## Supporting information

**S1 Fig. Test for locomotor defects in MBON candidates.** Single *MBON split-Gal4, UAS-CsChrimson* flies were placed in a circular agar bowl arena. Fly position was tracked under IR illumination with a camera, and then subsequently analyzed using ctrax and custom matlab scripts. The stimulation protocol was 3× (60s off, 30s on pulsing 633nm light at 50 Hz), and a total of 5 minutes of video was recorded for each trial. A) Velocity heat map, each row is an individual fly. Genotypes are denoted. B) Box and whiskers plot of average velocity over 3 trials, for light off and light on periods. Each data point is one fly, whiskers = 10th to 90th percentile, box = 25th to 75th percentile, and line in box = median. Statistical significance was calculated using unpaired Wilcoxon Rank Sum tests with Bonferroni correction, *p<0.05, **p<0.01, ***p<0.001. C) Bounded line plots. For each fly, the average velocity over 3 trials of (60s Light Off, 30s Light On) was calculated. The black line represents the mean average trial velocities of n = 16-20 flies for each genotype; the shaded grey areas represent the standard error. *MBON split-Gal4* lines crossed with *UAS-CsChrimson* are on top, paired with the genetic controls of *MBON split-Gal4* lines crossed with *w*^*1118*^, bottom. The red line marks the beginning of the light on period at 60s.

**S2 Fig. The SEZ neuron labeled by MB296B causes PER.** A) *MB296B split-Gal4* was crossed to *UAS-Chrimson88* and *UAS-CsChrimson* for light induced activation, and tested for proboscis extension in 3 conditions: (1) red light alone, (2) 30 mM sucrose to the tarsi, and (3) simultaneous red light and sucrose presentation to the tarsi. Extension rates were compared between each condition in the same fly and between different fly genotypes of the same condition for genetic controls (n = 27-58 flies). Values represent mean ± SEM. Statistical significance was calculated using paired Wilcoxon Rank Sum tests (same flies, different conditions) or unpaired Wilcoxon Rank Sum tests (flies of different genotypes, same treatment condition) with Bonferroni correction, *p < 0.05, ***p < 0.001. Green bars represent flies given sucrose and red light. Grey bars represent flies given sucrose. B) MB296B was inhibited with Kir2.1 and PER to 30 mM sucrose on the legs was recorded. Silencing with Kir2.1 increased PER (n = 44-56, Mean ± SEM). Statistical significance was determined by Wilcoxon Rank Sum tests with Bonferroni correction, *p<0.05. C) Candidates were silenced with 20xgtACR1 and PER to 30 mM sucrose on the legs was recorded, in the absence and presence of green light. n = 47, mean ± SEM. Statistical significance was determined by a paired Wilcoxon test. D) Top: Projection pattern of MB296B. Bottom: Projection pattern of the SEZ neuron labeled by MB296B, as determined by an intersection between the Hox gene scr and MB296B. E) *MB296B-split-Gal4* was crossed to *20xUAS-dsFRT-CsChrimson.mVenus*; *LexAop-FLP*/*CyO*; *scr-LexA*/*TM2* flies for light induced activation, and tested for proboscis extension to light (n = 31.) Of these 31 flies, 7 were also tested for their responses to 30 mM sucrose to the tarsi, and simultaneous red light and sucrose presentation to the tarsi as well. Statistical significance was calculated using Wilcoxon Rank Sum tests, **p<0.01.

